# Protein Folding accompanied by disulfide bond formation drives glutenin polymerization into multidimensional gluten networks

**DOI:** 10.1101/2025.06.23.661211

**Authors:** Boyu Xie, Jiahui Fu, Jihui Gao, Yi Li, Zhongxin Liang, D. Thirumalai, Dong D. Yang

## Abstract

Wheat flour dough, which is routinely used to make bread, pasta and noodles, can be stretched several thousand-fold without rupture. The molecular basis of this extraordinary property arises due to the ability to crosslink glutenin subunits, the major constituent of dough, into linear and branched elastic networks. The mechanism by which the glutenin subunits self-assemble and polymerize is unknown. Here, we show, using mass spectrometry, confocal imaging and molecular dynamics simulations that structuring of two hydrophobic residues, phenylalanine and tyrosine, in the core region of the monomeric glutenin 1Dx5 N-terminal domain (1Dx5-NTD) initiates inter molecular interactions and network formation. Upon folding of monomeric 1Dx5-NTD, two cysteine residues (Cys10 and Cys40) form an *intramolecular* disulfide bond and poises the third cysteine (Cys25) to engage in *inter molecular* crosslink with other glutenin subunits. Propagation of such a disulfide linkage pattern results in the formation of a cohesive linear gluten network. In alternate pathways, crosslinking of Cys10 with Cys25 with a third glutenin subunit results in the creation of a junction, which drives the formation of a three-dimensional network. The disulfide patterns in the networks accord well with the measured chemical reactivity of each cysteine residue and the solvent accessibility of the associated side chains during 1Dx5-NTD interactions. Our study shows that the diversity in the folding of a single glutenin subunit that exposes the third cysteine to the solvent is the key event that initiates polymerization and directs the formation of networks with differing architecture.

**Significance statement:** Glutenin proteins polymerize to form intricate molecular networks that support the expansion of dough by several thousand-fold. Although network formation occurs spontaneously, the polymerization mechanism is unknown. There are multiple cysteine residues in glutenin that can form intra and intermolecular disulfide bonds. Multiple experimental techniques and molecular dynamics simulations are used to show that folding of glutenin, initiated by two hydrophobic residues, exposes cysteines that are poised to form intermolecular disulfide bonds. The disulfide crosslinking pattern, determined by the initial folding process, dictates the creation of a linear or branched network. The study demonstrates that the propensity to form linear or branched networks is encoded by the diversity of the disulfide patterns created at the level of monomeric folding.

## Introduction

Elastic proteins, such as resilin, elastin, abductin and spider dragline silks, which exhibit high resilience, resist large strains and have low stiffness, are characteristics of rubber-like proteins(1). Despite their extraordinary mechanical properties, compared to synthetic polymers, little is known about how the monomers are cross-linked to form an elastomeric protein(2). Gluten has been part of the wheat flour-based food that endows substantial extensibility to dough and is routinely used to produce bread, noodles and pasta since the Upper Palaeolithic age(3). The major component of gluten is one of the largest protein molecule in nature, which is a glutenin polymer linked by disulfide bonds(4). Endogenous enzymes, such as wheat protein disulfide isomerase (PDI), catalyzes the formation of the gluten matrix during both the wheat growth and the bread making processes(5, 6). Exogenous molecules, such as sodium chloride, also impact the gluten network formation by altering glutenin conformations(7). Chemicals, usually oxidants, developed to impact the disulfide bonds formation are called dough rheology improvers, which are known to be hazardous to health(8).

As one of the storage proteins present in the starchy endosperm cells of wheat grain, glutenins are categorized as high molecular weight glutenin subunits (HMW-GSs) and low molecular weight glutenin subunits (LMW-GSs) (9). Though HMW-GSs represent only 8-10% protein in wheat grain, crosslinking between cysteine residues in HMW-GS is crucial for the gluten network formation (7, 9). Among allelically distinct HMW-GSs, 1Dx5 exhibits better dough processing properties than others(9, 10). We recently identified it as the major substrate protein of wheat PDI(5). It has long been recognized that the formation of intermolecular disulfide bonds is directly related to the rheological properties of dough while some of the disulfide linkages contribute more to dough rheology than others(6, 11). In noodles, the gluten network extends linearly in one dimension (1D) whereas in bread it extends in three-dimensions (3D)(12). We have previously showed that wheat PDI promotes the formation disulfide bonds in glutenin by favoring a β-strand structure without interfering with its folding(5, 13). However, it is unclear how glutenin folds into a structure that directs their subsequent polymerization and produces a polymer network of varying architecture in one and three dimensions(14).

Here, using multiple experimental techniques, such as mass spectrometry and confocal imaging complemented by atomic detailed simulations, we investigate how the glutenin subunits self-associate with each other to form specific linkage patterns to form 1D or 3D gluten network. We find that folding of hydrophobic amino acid residues in the protein core drives the association of HMW-GS 1Dx5 N-terminal domain (1Dx5-NTD) and poises a frustrated unpaired cysteine (C25) in 1Dx5-NTD to crosslink with cysteine residues in other glutenin subunits. Heterogeneous folding of a single molecule allows for different crosslinking patterns between 1Dx5-NTDs, resulting in branched protein skeleton formation and subsequent dough expansion in other dimensions. Our study produces a quantitative view of how an ensemble of proteins with diverse folding at the monomer level crosslink by distinct pathways to create networks with varying macroscopic properties. This work provides molecular insights into the mechanisms of self-assembly of proteins, which could be used to understand the formation of multidimensional extensible polymer networks in other systems such as filamentous actin and spider-silk (8, 12).

## Results

### Primary structure of 1Dx5-NTD and the gluten network

HMW-GSs contain a central repetitive domain flanked by an approximately 80-100 amino acid residue N-terminal domain (NTD) with 2 to 5 conserved cysteine residues and a 42 amino acid residue C-terminal domain (CTD) with 1 cysteine residue (Fig. 1a)(7). In HMW-GS 1Dx5, there are 3 cysteine residues in the N-terminal domain (1Dx5-NTD), 681 residues in the central repetitive region, and 1 cysteine residue in its CTD (*SI Appendix*, Fig. S1). Despite significant effort since almost seven decades ago, we have not determined the structure of HMW-GS 1Dx5(14). However, AlphaFold 3 predicts that it is largely unstructured, exhibiting characteristics of an intrinsically disordered proteins, in the central repeat domain while both the 1Dx5-NTD and CTD are partially folded(5, 15).

**Fig. 1.**
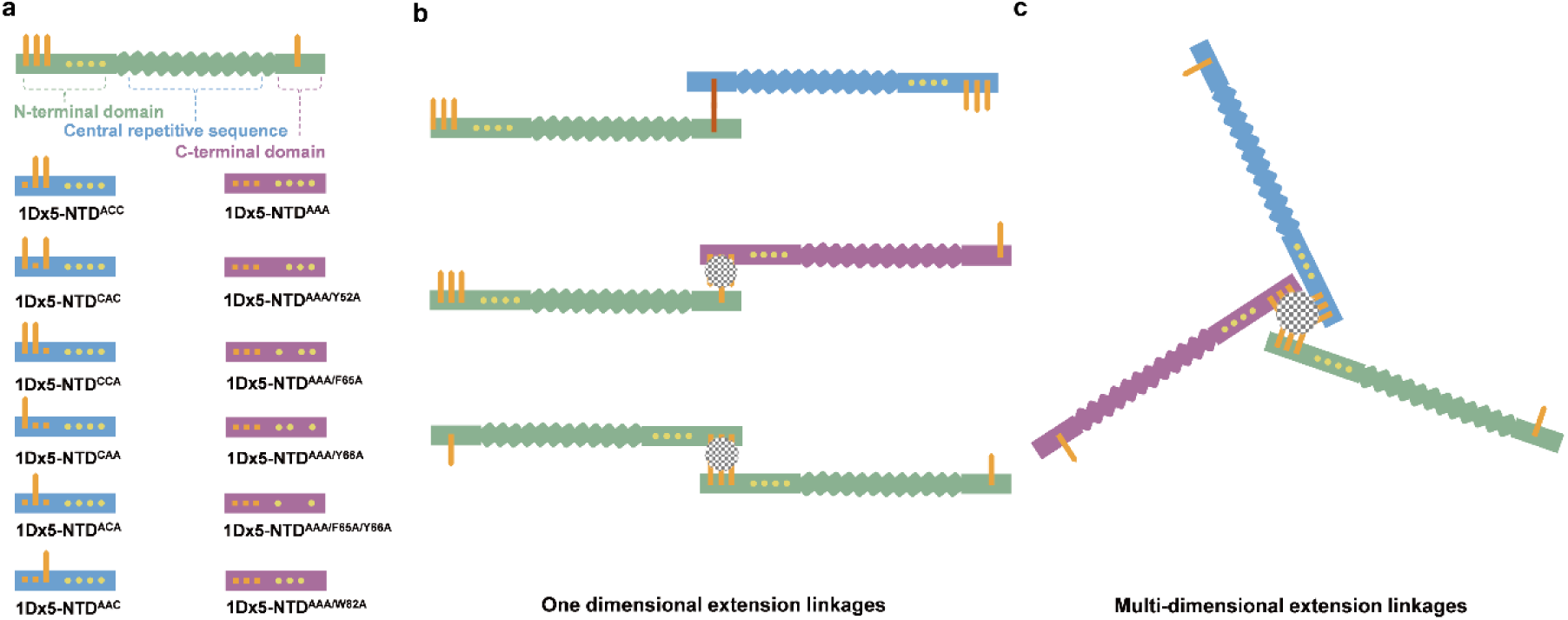
Model of glutenin 1Dx5 and their crosslinking. **a**, Structural model of high-molecular weight glutenin subunit 1Dx5. It contains an N-terminal domain (NTD) with 3 cysteine residues, a central repetitive domain and a C-terminal domain (CTD) with 1 cysteine residue. Unpaired C10, C25, C40 in the NTD and cystine residue in CTD are shown as orange sticks. Hydrophobic residues (Y52, F65, Y66, W82) are shown in yellow spheres. Blue ribbons are the variants of 1Dx5-NTD with one or two cysteine residues while the other cysteine residues are mutated to alanine. Purple ribbons are the cysteine free 1Dx5-NTD variants with mutations of hydrophobic residues Y52, F65, Y66, W82 are mutated to alanine, respectively. **b**, Crosslinks between glutenins can be decomposed into linear and branched crosslinking. Three possible ways of linear crosslinking, the tail-to-tail (green and blue ribbons), head-to-tail (green and purple ribbons), and head-to-head crosslinking (green and green ribbons). Paired disulfide bond is in red stick. **c**, Branched crosslinking occurs when a third glutenin participates in the polymerization at one joint and introduces multi-dimensional extension. Yellow dots represent hydrophobic residues on the N-terminal of a glutenin subunit. Mosaic circles represent disulfide linkages with unknown crosslinking pattern.

To form an 1D gluten network, glutenin subunits should be linked linearly in a either tail-to-tail, head-to-tail, or head-to-head fashion (Fig. 1b). For multidimensional gluten network formation, the crosslinking pattern should also support branched extension of the protein skeleton where more than two glutenin molecules crosslink at a single joint (Fig. 1c). There can only be one crosslinking pattern in the tail-to-tail arrangement, but it is unknown how disulfide bond formation in 1Dx5-NTD affects the geometry of a gluten network (mosaic regions in Fig. 1b and c). Prior to the formation of linear network, glutenin molecules must recognize each other and self-assemble by a process whose details are unknown. Here, we used mutations in the cysteine residues in 1Dx5-NTD^WT^ to generate variants in which two cystines residues are retained while the other is mutated to alanine, labeled as 1Dx5-NTD^ACC^, 1Dx5-NTD^CAC^, and 1Dx5-NTD^CCA^. We also created variants with only one cystine residue with the other two being mutated to alanine, labeled as 1Dx5-NTD^CAA^, 1Dx5-NTD^ACA^, and 1Dx5-NTD^AAC^. Finally, we also considered triple mutants with no cysteines residues as 1Dx5-NTD^AAA^ (Fig. 1a and *SI Appendix*, Table S1). The results of the mutant folding prove the essential first glimpse of the network formation.

### Folding directs 1Dx5-NTD association

1Dx5-NTD molecules must be in proximity to facilitate crosslinking to spontaneously associate. To exam the stability of 1Dx5-NTD associations, cystine free 1Dx5-NTD^AAA^ triple mutants were utilized to eliminate covalent crosslinking between cysteine residues that could interfere with the non-covalent interactions. We used isothermal titration calorimetry (ITC) to determine the thermodynamic changes during the dilution of 1Dx5-NTD^AAA^ associations(16). Results showed that 1Dx5-NTD^AAA^ association were quite stable that dilution caused no dissociation, as verified with different scale of pre-calibrated ITC instruments (*SI Appendix*, Fig. S2a-b).

We then used salting in effect to examine the protein association dynamics with sedimentation velocity analysis(17). At a rotor speed of 40k rpm, two major species of around 60 kDa and 100 kDa (corresponding to hexameric and decameric association) were found and their abundances were barely affected by salt concentrations (Fig. 2a-c). However, large particles of ∼36,000 kDa observed at lower rotor speed of 10k rpm decreased at 100 mM NaCl (*SI Appendix*, Fig. S2c-e). This was further examined by dynamic light scattering, and the size of 1Dx5-NTD^AAA^ association decreased from ∼500 nm to ∼200 nm when the salt concentration increased from 0.01 mM to 0.1 mM (Fig. 2d). These results suggested that association of 1Dx5-NTD was mainly driven by electrostatic interactions.

**Fig. 2.**
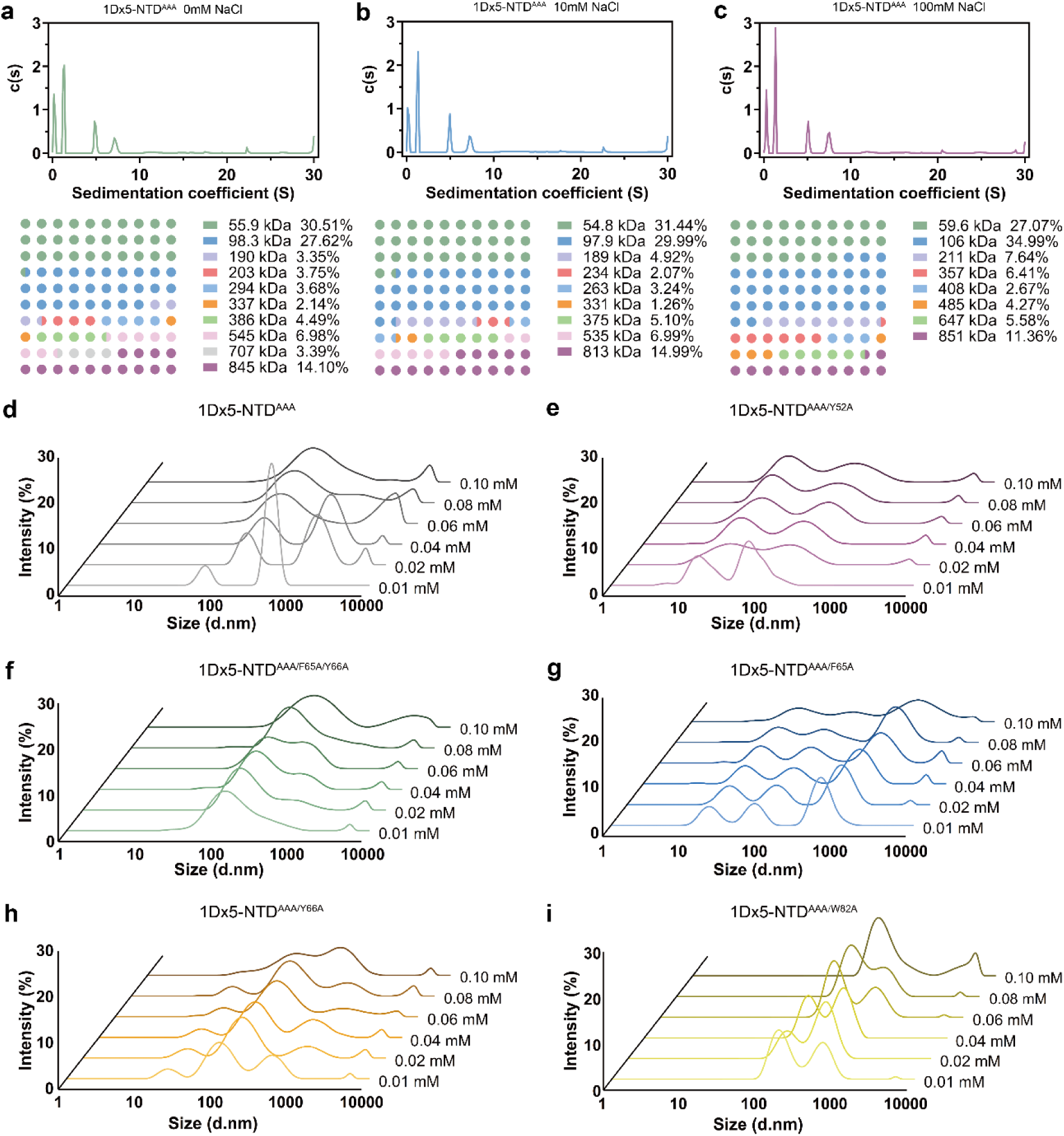
Role of hydrophobic residues in 1Dx5-NTD association. **a**-**c**, Sedimentation velocity profiles of cystine free 1Dx5-NTD^AAA^ protein at different NaCl concentrations (0mM, 10mM, and 100mM) and 40k rpm. Below each are the ratios of different molecular weight protein associations. **d**-**i**, Dynamic light scattering profiles of hydrophobic amino acid residue mutations on the 1Dx5-NTD^AAA^ protein association behavior at different NaCl concentrations (0.01mM, 0.02mM, 0.04mM, 0.06mM, 0.08mM, and 0.10mM).

On the other hand, it was initially assumed that 1Dx5-NTD association was driven by interactions between hydrophobic amino acid residues. To test this, we mutated four of these residues (yellow dots, Fig. 1a) to assess their significance in the 1Dx5-NTD association process. Mutation Y52A and double mutant, F65A/Y66A in 1Dx5-NTD^AAA^ shifted the assembly sizes to ∼90 nm and ∼120 nm, respectively, even at low salt concentration of 0.01 mM (Fig. 2e-f). Mutations F65A, Y66A and W82A also decreased the 1Dx5-NTD^AAA^ assembly size at low salt concentrations (Fig. 2g-i). These results indicated significant roles these hydrophobic residues played in 1Dx5-NTD molecular associations.

It was puzzling that the 1Dx5-NTD association was mainly driven by electrostatic interactions while these hydrophobic residues were vital to their integrity. To resolve this apparent contradiction, we used molecular dynamics (MD) simulations to picture the processes of 1Dx5-NTD molecular folding and interactions. First, folding of a single molecule was performed (*SI Appendix*, Fig. S3a). 1Dx5-NTD exhibited overall disordered architecture, with Y52, F65, Y66, and W82 forming the core region (Fig. 3a), as verified by their decreased solvent accessible surface area (SASA, *SI Appendix*, Fig. S3b-e). Next, the process of two or three 1Dx5-NTD molecules interacting with each other were simulated (*SI Appendix*, Fig. S3a). Mild to moderate structural adjustments were observed between 1Dx5-NTD molecules interacting with another (*SI Appendix*, Fig. S3f-g) or two other molecules (*SI Appendix*, Fig. S3h-j), as suggested by their structural deviation with root-mean-square deviation (RMSD) ranging from 3.425Å to 9.809Å. The large variation in the RMSD suggests that there is structural heterogeneity during the 1Dx5-NTD association process. Consistently, these four hydrophobic residues were all structured in the hydrophobic core region of each 1Dx5-NTD molecule in all interaction scenarios (Fig. 3b-c).

**Fig. 3.**
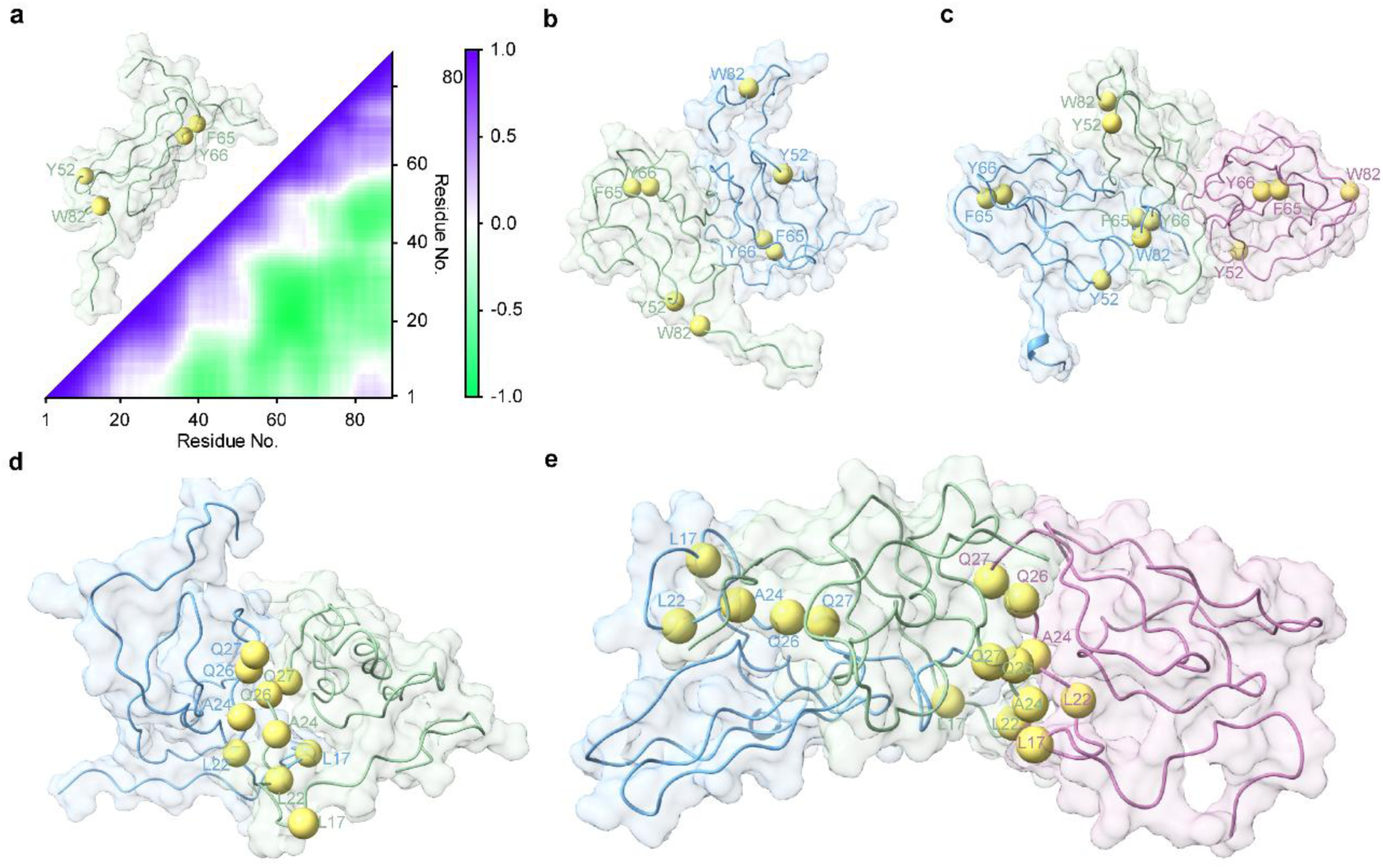
Role of hydrophobic residues in 1Dx5-NTD folding. **a**, Dynamic map of correlated motions during the folding of one 1Dx5-NTD molecule calculated from the MD simulation. The folded structure is shown in the upper triangle and key hydrophobic residues are shown in yellow spheres. **b**, Conformation of two 1Dx5-NTD molecules interacting with each other. One of the molecules is colored in green and the other is colored in blue. Key hydrophobic residues (yellow spheres) folded into the core are labelled in green and blue, respectively. **c**, Conformation of three 1Dx5-NTD molecules interacting with each other. They are colored in green, blue, and purple, respectively. Key hydrophobic residues (yellow spheres) folded into the core are labelled in green, blue, and purple, respectively. **d**, Interface of two 1Dx5-NTD molecules interacting with each other. C_α_ of key residues involved in protein interaction (yellow spheres) are labelled in green and blue, respectively. **e**, Interface of three 1Dx5-NTD molecules interacting with each other. Key residues involved in protein interaction (yellow spheres) are labelled in green, blue and purple, respectively.

Analysis of the intermolecular interaction between two 1Dx5-NTD molecules found that A24 in both the molecules was involved in hydrogen bonding and hydrophobic interactions (Fig. 3d and *SI Appendix*, Fig. S4a). L22 and Q27 in both the molecules participated in the hydrophobic interactions between two 1Dx5-NTD molecules (molecule A and B). In the 1Dx5-NTD trimer interaction, A24 in all three molecules, L22 in molecules A and B, L17 and Q26 in molecules B and C participated in hydrophobic interactions (Fig. 3e and *SI Appendix*, Fig. S4b). Cross-correlation analysis showed that structuring in above mentioned five residues were highly negatively correlated with residues F65 and Y66 during 1Dx5-NTD folding with correlation <−0.75 (Fig. 3a). Thus, it is the folding of hydrophobic residues into the protein core that poises the electrostatic interactions between L17, L22, A24, Q26 and Q27 in 1Dx5-NTDs.

### Role of cystine residues in gluten network formation

To identify the role of each cystine residue in the gluten network formation, 1Dx5-NTD^WT^ and its mutant proteins with one retained cysteine residue were purified, added to dough with approximately 10%, 25%, 50% and 100% of the total content of HMW-GS N-terminal, respectively(9). Dough with different protein addition was stained with Rhodamine B, imaged under confocal microscope and the gluten network was quantitatively analyzed(5, 12). The protein percentage area increased significantly, from 32.2% to 41.5%, as 1Dx5-NTD^WT^ protein addition increased from 10% to 100% (Fig. 4a and *SI Appendix*, Fig. S5a). Addition of 1Dx5-NTD^CAA^ and 1Dx5-NTD^AAC^ also increased the protein percentage area to a maximum of 39.0% and 39.8%, respectively. Increasing the amount of 1Dx5-NTD^ACA^ to approximately 50% enhanced the gluten network protein percentage area to 39.4%. However, further addition of 1Dx5-NTD^ACA^ decreased the protein percentage area to 37.9%. The values of the junction density and total protein length were enhanced upon addition of 1Dx5-NTD^WT^ from 10% to 100% (Fig. 4b-c and *SI Appendix*, Fig. S5b-d). Similarly, addition of 1Dx5-NTD^CAA^ and 1Dx5-NTD^AAC^ also increased these values. In contrast, addition of 1Dx5-NTD^ACA^ increased these values first and then decreased after 50% addition. These results suggested that C25 in 1Dx5-NTD contributed differently from C10 and C40 in the gluten network formation.

**Fig. 4.**
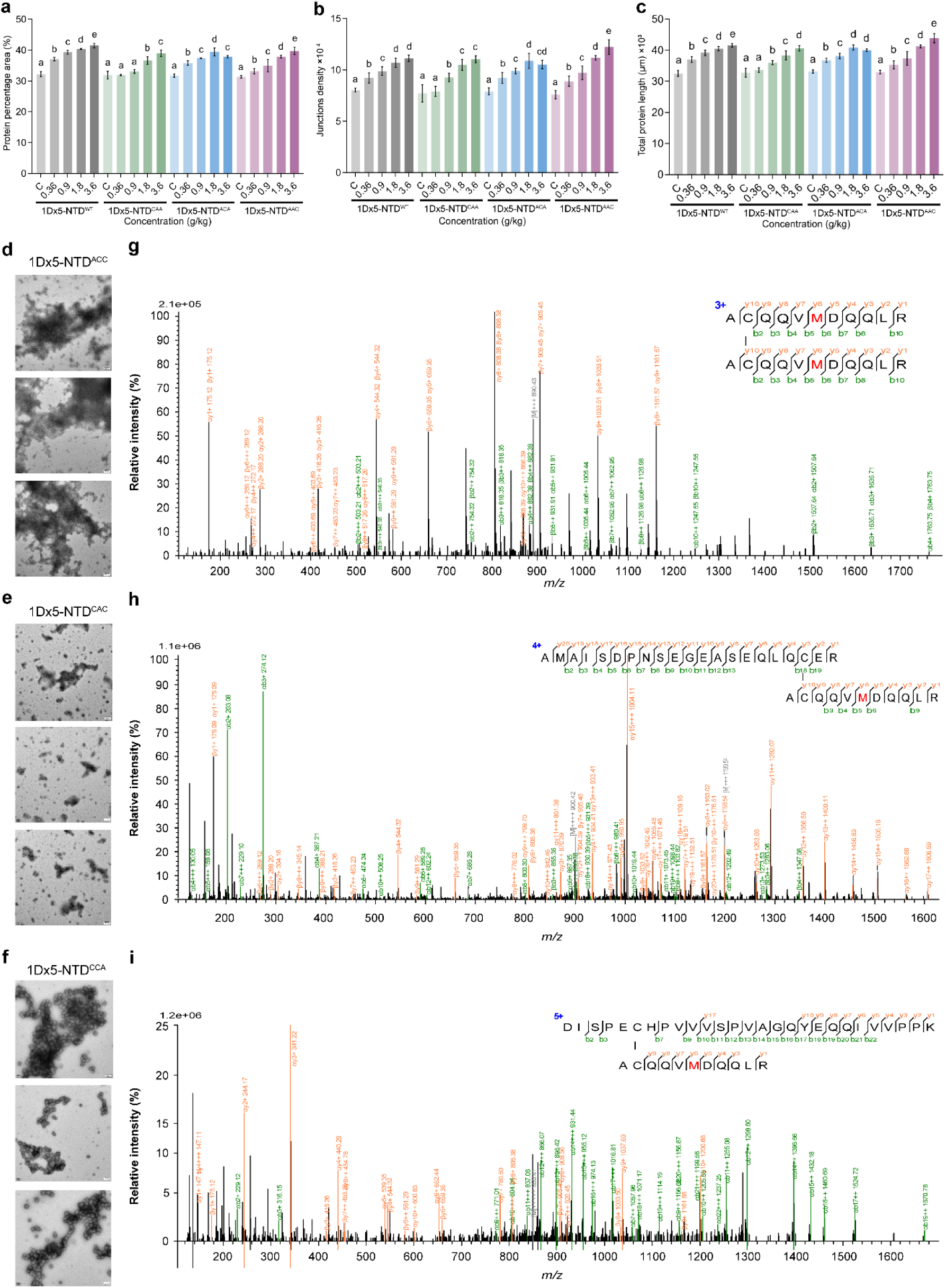
Role of cystine residues in gluten network formation. **a**-**c,** Quantitative analysis of the gluten network to get the protein percentage area, junction density and total protein length of dough gluten network formed with different 1Dx5-NTD variant addition. Dough was made with addition of 1Dx5-NTD^WT^ protein and its cystine residue mutants corresponding roughly to 10%, 25%, 50%, 100% of the total high molecular weight glutenin subunits NTDs. 1Dx5-NTD^CAA^ represents C25A/C40A mutation, 1Dx5-NTD^ACA^ represents C10A/C40A mutation, and 1Dx5-NTD^AAC^ represents C10A/C25A mutation. Different letters indicate significant analysis of differences at *p*<0.05. **d**-**f**, Electron microscopic images of gluten network formation by 1Dx5-NTD cystine mutant proteins with two cysteine residues maintained while C10, C25 and C40 were mutated into alanine, respectively. 1Dx5-NTD^ACC^ represents C10A mutation, 1Dx5-NTD^CAC^ represents C25A mutation, and 1Dx5-NTD^CCA^ represents C40A mutation. Scale bars represent 1 μm. **g**-**i**, Annotated MS/MS spectrums of the digestion product of the crosslinking products of two 1Dx5-NTD^WT^ molecules, and the precursor charge and m/z are shown in the spectrums, indicating C25-C25, C10-C25, C40-C25 disulfide bonds, respectively.

To further identify the role of each cystine residue during gluten network formation, 1Dx5-NTD mutant proteins in which two cystine residues maintained were purified and subjected to crosslinking studies. These variants were capable of not only association but also polymerization. Under electron microscope, we found that 1Dx5-NTD^ACC^ and 1Dx5-NTD^CCA^ formed continuously cohesive protein network while 1Dx5-NTD^CAC^ only formed a small and discontinues protein network (Fig. 4d-f). It follows that C25 was the key to propagate the protein skeleton while C10 and C40 formed a gluten network of limited-size.

To identify the pairing patterns between cystine residues during gluten network formation, 1Dx5-NTD^WT^ protein on which all three native cystine residues present were subjected to crosslinking studies. We then isolated the reaction products of two 1Dx5-NTD molecules crosslinked together in non-reducing SDS-PAGE, in-gel digested, and the cystine crosslinks were analyzed with MS/MS (*SI Appendix*, Fig. S6a). Annotated MS/MS spectra indicated the formation of C25-C25 (Fig. 4g), C10-C25 (Fig. 4h), and C40-C25 (Fig. 4i) disulfide linkages between two 1Dx5-NTD molecules. These results again clearly illustrate that C25 was always involved in the intermolecular crosslinking while C10 and C40 were involved in the formation of auxiliary disulfide linkages.

### Folding directed 1Dx5-NTD crosslinking

We used MD simulations to image the disulfide bonds formation process during 1Dx5-NTD folding and the associated interactions with other molecules. In the folding process of single 1Dx5-NTD molecule, the distance between the Cα of C10 and C40 reached approximately 9 Å, which is sufficient for disulfide crosslinking to take place (Fig. 5a). In contrast, intramolecular distance between Cα of C25 and C10 (or C40) never reached close enough distance for disulfide bond formation. For two molecules interacting with each other, C10 and C40 in each 1Dx5-NTD formed intramolecular disulfide bonds while C25 residues reached within 9 Å for intermolecular crosslinking (Fig. 5b). Distances between Cα of all other pairs of cystine residues were beyond 9 Å (*SI Appendix*, Fig. S6b). When three molecules interact with each other, C10 and C40 in two 1Dx5-NTD molecules formed intramolecular disulfide bonds (molecules A and C, Fig. 5c). Strikingly, C25 in one of the 1Dx5-NTD molecules (molecule A) formed intermolecular linkage with C25 in the 2^nd^ molecule (molecule B, Fig. 5c) and C25 in the 3^rd^ 1Dx5-NTD molecule (molecule C) crosslinked with C10 in the 2^nd^ molecule (molecules B, Fig. 5c). The Cα distances of other cystine residues were not close enough for disulfide bond formation (*SI Appendix*, Fig. S6c).

**Fig. 5.**
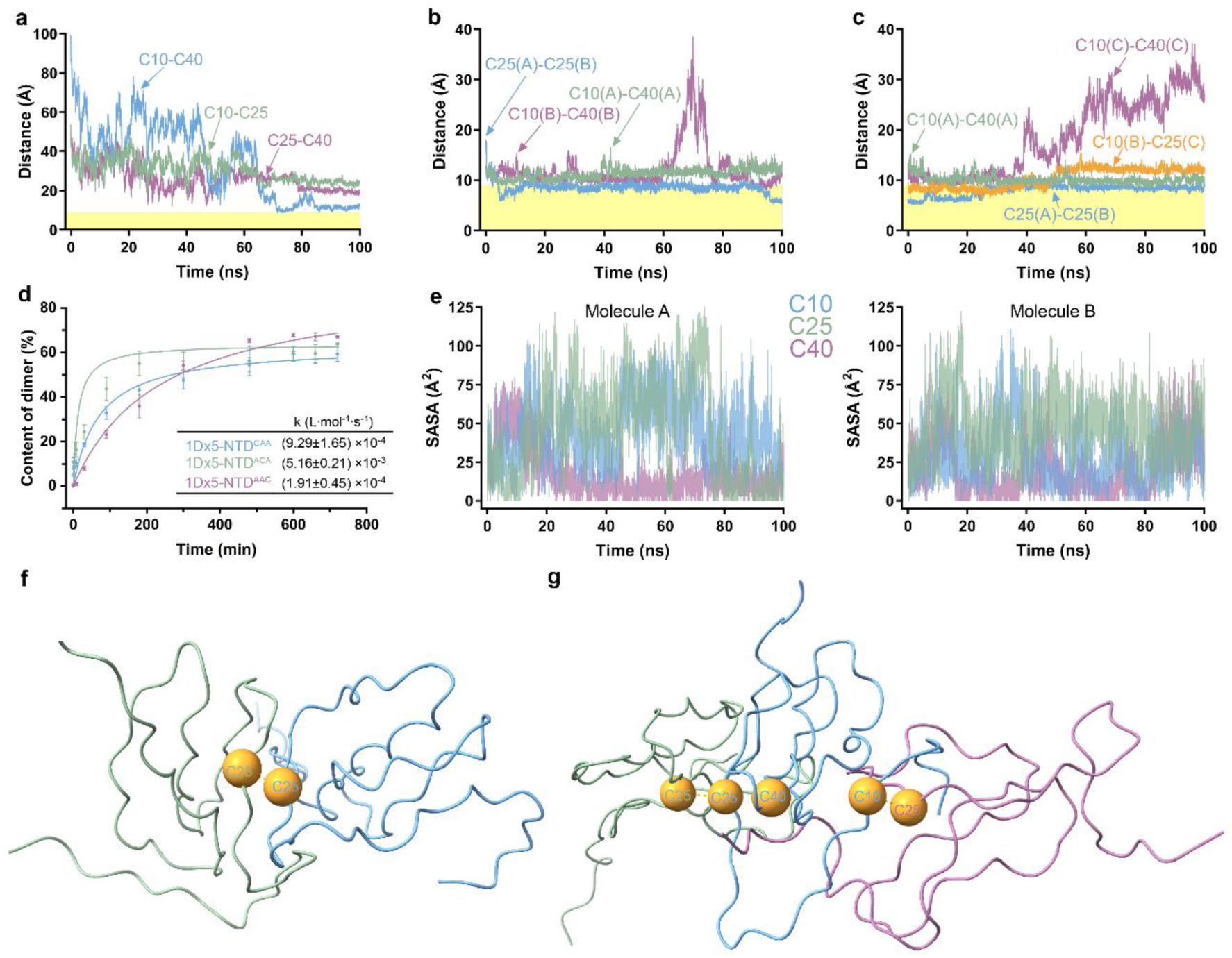
Folding drives disulfide formation for linear and branched linkages. **a**-**c**, Distances between C_α_ of different cystines during one 1Dx5-NTD molecule folding, two 1Dx5-NTD molecules interaction, and three 1Dx5-NTD molecules interaction, respectively. **d**, Self-crosslinking kinetics of 1Dx5-NTD cystine mutant proteins. 1Dx5-NTD^CAA^ represents C25A/C40A mutation, 1Dx5-NTD^ACA^ represents C10A/C40A mutation, and 1Dx5-NTD^AAC^ represents C10A/C25A mutation. The crosslinking products were quantified and each reaction rate is listed in the table inserted. **e**, Solvent accessible surface area (SASA) of the side chain in each cystine residue during two 1Dx5-NTD molecules (molecules A and B) interacting with each other, respectively. **f**, Conformation of two 1Dx5-NTD molecules interacting with each other while the distance between cystine residues for intermolecular disulfide bond formation is allowed. Green and blue cartoon represent two molecules and orange spheres represent cystine residue. **g**, Conformation of three 1Dx5-NTD molecules interacting with each other. Green, blue and purple cartoon represent three molecules and other symbols are the same as in f.

To measure the chemical properties of the cysteine residues that lead to such pairing pattern during 1Dx5-NTD polymerization, protein variants 1Dx5-NTD^CAA^, 1Dx5-NTD^ACA^ and 1Dx5-NTD^AAC^ consisting of one cystine residue were subjected to crosslinking rate determination. These mutations did not significantly alter the secondary structure content or structural stability of 1Dx5-NTD (*SI Appendix*, Fig. S7a-b). Crosslinking products were analyzed in non-reducing SDS-PAGE and quantified (*SI Appendix*, Fig. S6d-f). C25 exhibited the maximal reaction rate among all three cystines, which was ∼5.6 fold greater than C10 and ∼27 folds of C40 (Fig. 5d). SASA of each cystine residue in 1Dx5-NTD during two molecules interaction were analyzed as a proxy for reactivity(18). We found C25 in molecule A exhibited the highest SASA during most of the simulation time, slightly exceeding C10 and much higher than C40 (Fig. 5e). This readily explained the observation that 1Dx5-NTD^ACA^ exhibited the highest self-crosslinking reaction rate followed by 1Dx5-NTD^CAA^ and then 1Dx5-NTD^AAC^. Meanwhile, C10 in molecule B has the highest SASA most of the time, followed by C25 and C40, which explains the crosslinking product between 1Dx5-NTD^WT^ where C25-C10, C25-C25 and C25-C40 were all observed (19). In general, C25 in one molecule was always involved in the crosslinking between C10, C25, and C40 in the other molecule.

Interestingly, structuring of F65 and Y66 were also highly negatively correlated with C25 during folding of 1Dx5-NTD with correlation <−0.75, while moderately negative with C10 with correlation between −0.75 and −0.5 (Fig. 3a). C40, on the other hand, was not significantly correlated with these two hydrophobic residues. Thus, the molecular architecture induced by folding of the hydrophobic residues positions C25 at the bimolecular and trimolecular interface for C25-C25 disulfide bond formation (Fig. 5f-g). This is the pattern conducive to 1Dx5-NTD linear polymerization needed for one-dimensional gluten network extension (Fig. 6a). When three 1Dx5-NTD molecules interact, C25 was covalently linked to one molecule and C10 also participated in the intermolecular crosslinking with the third molecule (Fig. 6a and b). Such motif is the molecular foundation of protein skeleton branching for multi-dimensional extension of the gluten network (Fig. 6b). It is remarkable that the formation of 1D or 3D network is determined solely by the pattern of intermolecular disulfide formation involving a few protein molecules. Intermolecular disulfide bond formation involving only C25 results in the formation of 1D whereas disulfide bond involving C10 and C25 produces 3D network (Fig. 6c).

**Fig. 6.**
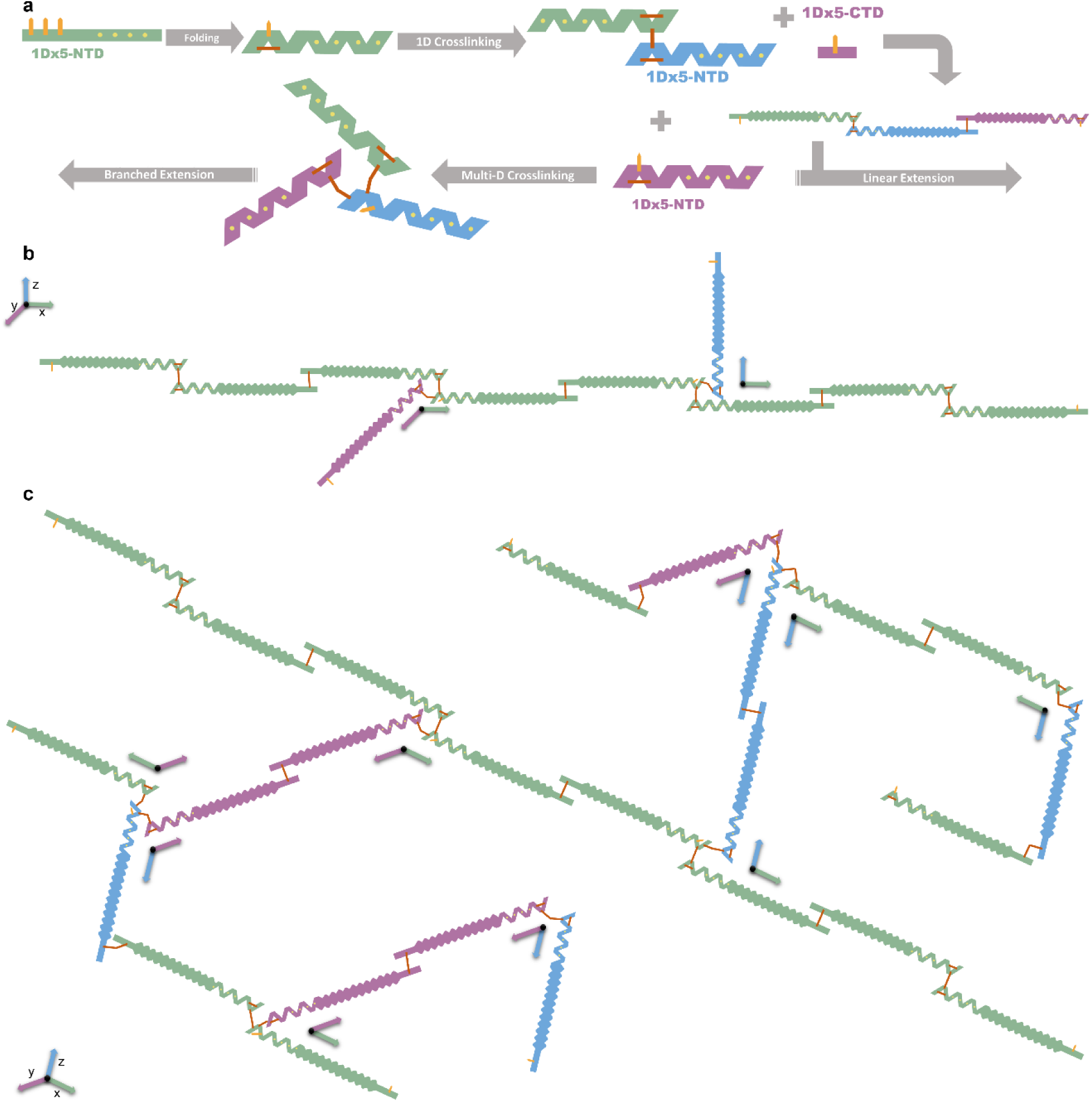
Disulfide patterns in the 1D and 3D gluten network. **a**, Cartoon representation of 1Dx5 polymerization. 1Dx5-NTD folds and forms intramolecular bonds that allows intermolecular crosslinking between C25 residues. Starting from this joint, crosslinking with another 1Dx5-CTD allows elongation in 1D while crosslinking with another 1Dx5-NTD joint elongates the protein skeleton in another dimension. Frustrated unpaired cysteine residues are shown in orange sticks and paired disulfide bonds are shown in red sticks. **b**, Example of crosslinking of a third 1Dx5-NTD to one joint of the linearly polymerized glutenins (green on x-axis) introduces extension to the y-axis (purple) or the x-axis (blue). **c**, Propagation of such linear and/or branched crosslinking forms a gluten network that allows extension on multi-dimensions.

## Discussion

Using a range of experimental techniques and molecular dynamics simulations, we have discovered the distinct ways in which the folding of monomeric glutenin drives complex polymerization reactions, thus creating intricate cohesive protein networks with varying morphology in dough. Glutenins have to associate with each other for the cysteine residues to crosslink by forming disulfide bonds(20). For this to occur, glutenin monomer must fold, which is driven by hydrophobic interactions that brings the cysteine residues in proximity to form disulfide bonds. Upon folding, multiple glutenin subunits associate with each other in solution and is not disrupted even upon dilution(16), which implies that assembled state is stable. Interestingly, folding is initiated through hydrophobic interactions facilitate glutenin association, as it is the case in other proteins that in which disulfide bond formation plays an important role(21, 22). For instance, single molecule pulling experiments in an engineered immunoglobin with a single disulfide bond showed that the bond between the cysteines forms only after the protein folds by hydrophobic interactions(23). This finding is also borne out in glutenin in which the central repetitive domain with hydrophobic residues play significant roles during the network formation(20). In particular, the hydrophobic residues Y52, F65, Y66 and W82 are directly involved in the intermolecular interactions that lead to 1Dx5-NTD association. Indeed, mutation of these residues decreased the size of molecular association.

### Central role of Cysteine 25

There are three cystine residues in 1Dx5-NTD. However, it is unknown if the disulfide bonds form randomly or if there are specific crosslinking patterns that confer multi-dimensional extensibility to the polymer network. The gluten network analysis presented here, shows that C25 plays a different role in the gluten network formation compared to the others. First, MS/MS analysis of 1Dx5-NTD crosslinking products shows that C25-C25, C10-C25 and C40-C25 disulfide linkage patterns are present, suggesting that C25 is always involved in 1Dx5-NTD polymerization reaction. Second, EM images show that C25 is necessary to form a cohesive gluten network. These findings are in accord with the crosslinking rate assay of each cysteine residue which shows that C25 has the highest reaction rate. It is also well explained by MD simulations, which show that folding of hydrophobic residues F65 and Y66 is necessary to bring C10 and C40 in proximity to form an intramolecular disulfide bond in the monomeric 1Dx5-NTD. Upon folding of the hydrophobic core C25, with the largest SASA, is poised for inter-molecular crosslink formation, resulting in network formation. The finding that 1Dx5-NTD variants bearing only C10 or C40 also crosslinks together confirms the capability of C10 and C40 in the formation of intermolecular disulfide linkages, but at a slower rate. Our results are consistent with previous findings that folding of the central core to the native state is necessary for disulfide bond formation(13, 24).

When more than two 1Dx5-NTD molecules are involved in the crosslinking reaction, the C10-C40 intramolecular disulfide bond in one of them is not formed, leaving C10 accessible for intermolecular crosslinking with the third molecule. This is consistent with the crosslinking rate assay that the reaction rate of C25 is followed by C10 and then C40, as well as MD simulation results which show that C10 has the largest SASA in the other interacting 1Dx5-NTD molecule followed by C40. And this offers auxiliary disulfide linkages C25-C10 and C25-C40, which makes branching of the protein skeleton in a gluten network possible. It is worth mentioning that although other crosslinking patterns were not observed in the simulation, they could still be present in reality due to spontaneous or enzyme catalyzed disulfide exchange that may produce crosslinking pattern different from the initial reaction product(25).

In summary, we have established that multi-dimensional extension of dough is a result of the formation of a gluten network, which occurs in steps. First, folding of monomeric glutenin initiates the polymerization reaction. Folding of 1Dx5-NTD is driven by the hydrophobic clustering of aromatic amino acid residues F65 and Y66 in the core, which directs not only residues that are poised for 1Dx5-NTD interactions, but also renders the proximity of C25 in one molecule and C25, C10 or C40 in another 1Dx5-NTD for their subsequent crosslinking. The main disulfide linkage is C25-C25 for linear protein polymerization and C25-C10 while C25-C40 disulfide linkages are needed for auxiliary reactions to generate branched elongation of a glutenin polymer. Our results provide a detailed mechanism of the folding-directed association and crosslinking of glutenins into a proteinaceous polymer that is routinely used in the food industry.

## Materials and methods

### Mutation, expression, and purification of 1Dx5-NTD and its variant proteins

Plasmid pET-30b harboring gene encoding 1Dx5-NTD was kindly provided by Prof. Songqing Hu (South China University of Technology) and was transformed into *E. coli* BL21 (DE3) cells (Tsingke Biotechnology Co., Ltd., Beijing, China). 1Dx5-NTD mutants were constructed with primers and PCR conditions as described in Table S1 in the Supporting Information. Each protein expression was induced by culturing *E. coli* cell harboring the corresponding plasmid in Luria-Bertani medium with kanamycin till OD_600_ reached 0.6-0.8 before the addition of a final concentration of 1mM isopropyl β-D-thiogalactoside. Cells were disrupted by sonication, cell debris was removed by centrifugation, and the supernatant was passed through a pre-equilibrated Ni^2+^ column (QIAGEN, Germany). Protein was eluted with buffer containing increasing concentration of imidazole, and further purified with gel filtration chromatography equipped with column Superdex 75 10/300 GL (Cytiva, US).

### Isothermal titration calorimetry

On an MicroCal VP-iTC or iTC_200_ calorimeter (Microcal Inc., Northampton, MA, USA), 50 mM PBS buffer pH 7.4 was titrated into 0.23 mM 1Dx5-NTD^AAA^ triple mutant protein solution with 20 successive additions of 10 μL PBS buffer and interval time of 210s for VP-iTC, or 2 μL PBS buffer with interval time of 120s for iTC_200_. Thermographs were integrated with the ORIGIN software (Microcal Inc.) by plotting heat change against the addition of PBS into the protein.

### Sedimentation velocity

In an XL-I analytical ultracentrifuge (Beckman Coulter, Fullerton, CA, USA) equipped with an AnTi60 rotor at 20℃, approximately 400 μL 1Dx5-NTD samples or the reference buffer to which the protein was thoroughly dialyzed against were added to double-sector centerpieces. Radial absorbance at 280 nm was recorded at 5 min intervals with radial increment of 0.002 cm in continuous scanning mode and rotor speed of 40k rpm and 10k rpm, respectively. Molecular mass distribution *c*(*M*) was obtained by fitting the sedimentation velocity data with molecular weights ranging from 54.8 to 851 kDa and 259 to 37800 kDa, respectively. The friction coefficient was varied during the fitting procedure to obtain the best-fit value.

### Dynamic light scattering

1Dx5-NTD^AAA^ and its variants were diluted to the concentration of 0.01 mM, 0.02 mM, 0.04 mM, 0.06 mM, 0.08 mM and 0.10 mM, and placed in transparent quartz cuvettes in a temperature-controlled thermal chamber of a Zetasizer Nano ZEN3700 instrument (Malvern Instruments, Malvern, UK) at 25℃. The protein particle size distribution was analyzed with average of three replicates.

### All-atom Molecular Dynamics simulations

CHARMM36 force field was used to perform atomically detailed molecular dynamics simulations of 1Dx5-NTD, including one 1Dx5-NTD molecule alone, two 1Dx5-NTD molecules interaction, and three 1Dx5-NTD molecules interaction, respectively(26). Sodium ions were used to neutralize the system, which was explicitly solvated using the TIP3P water potential in a box with a minimum clearance of 8 Å between solute and the side of the box(27). Particle Mesh Ewald molecular dynamics (PMEMD) CUDA version was employed to run the molecular dynamics simulation on GPUs with the GROMACS 2020.3 package(28, 29). We first performed energy minimization for 5000 steps to repair the asymmetrical geometries in the system with the determined force of 1000.0 kJ/mol/nm. Particle Mesh Ewald (PME) method was used for long-ranged electrostatic interactions whereas the geometry of bond length was confined using the Steepest Descent Algorithm (SDA) method. The NVT (temperature) and NPT (pressure) systems were fixed at 300 K and 1.0 bar after energy minimization using the Parrinello-Rahman barostat method, water molecule geometry was restrained using the SETTLE algorithm, and the non-water bonds were restrained using the LINCS algorithm. Finally, an unrestrained 100 ns run was carried out under the same conditions as in the equilibrium step. The snapshots were saved every 10 ps. Built-in modules in GROMACS were used to analyze the data(30, 31).

### Confocal microscopy and gluten network analysis

Dough containing different concentrations of 1Dx5-NTD proteins, wild type or mutants, were prepared as described previously(7, 12). The dough sample was cut and transferred to an object carrier, sealed with a cover glass, and observed with an Olympus IX81 inverted microscopy equipped with a FluoView FV1000 confocal system (Olympus, Tokyo, Japan) and a 60×objective lens (UPlanSApo 60×, NA=1.35, WD=0.17). The excitation wavelength was set at 561 nm and the emission wavelength at 405/633 nm was detected. Different spots on the x-y-axis were recorded for each sample and dough samples were produced 5 times. The gluten network in each confocal microscopic image was analyzed with the AngioTool64 v.0.6a (National Cancer Institute, NIH, MD, USA)(12).

### Transmission electron microscopy

Protein variant was reduced with fresh dithiothreitol (DTT) before desalted with PD-10 column (GE health care, Chicago, USA). The protein concentration was adjusted to 5.0 mg/mL and then incubated in buffer with 1 mM KBrO_3_ at 4 ℃ for 12 h. 3 μL of each sample was loaded on the grid for 30 s and then stained with 0.5% uranyl acetate for 30 s. The specimen was imaged on a Tecnai Spirit electron microscope (FEI^TM^) operated at 100 kV equipped with a CCD camera (FEI^TM^ Eagle) and images were collected at a nominal magnification of 30000×.

### Mass spectrometry assisted identification of cross-links

1Dx5-NTD^WT^ was allowed to crosslink under the oxidation conditions for a set of time intervals and the crosslinking product was applied to gel electrophoresis. The protein band on the gel corresponding to two 1Dx5-NTD molecules crosslinking products was cut and digested with trypsin overnight, and the digestion product was eluted with gradients of acetonitrile. On a nanoLC-Q Exactive HF Orbitrap mass spectrometry (Thermo Fisher Scientific, Waltham, Massachusetts, U.S.), digested peptides were loaded to an analytic reverse phase C_18_ (75 μm×20 cm, 3 μm) column with a linear gradient from 92% buffer A (0.1% formic acid in water) to 22% buffer B (0.1% formic acid in acetonitrile) in 50 min, and then 78% buffer A to 32% buffer B in 12 min at a flow rate of 300 nL/min. The top 20 most intensive precursor ions from each full scan (300-1600 m/z) were isolated for HCD MS^2^ with a dynamic exclusion time of 40 s. Peptides were identified with the Proteome Discoverer (1.4.0.288) and the crosslinks between cysteines were identified with pLink 2.3.7 where the precursor mass accuracy was set at 10 ppm, the fragment ion mass was set at 20 mDa, and the results were filtered by applying a 5% FDR cutoff at the spectral level(32).

### Circular Dichroism

1Dx5-NTD protein concentration were adjusted to 0.2 mg/mL and placed into a quartz cuvette with 1 cm path length. On a Chirascan circular dichroism polarimeter (Applied Photophysics, U.K.), CD spectrum was recorded by scanning the sample from 190 nm to 260 nm at 25℃ with a bandwidth of 1 nm and step size of 1 nm. Secondary structural content was estimated with the built-in CDNN software. To determine the melting point of 1Dx5-NTD and its variants, protein sample in the thermostatic chamber was heated from 20℃ to 105℃ and CD signal at 222 nm was recorded. Fraction of unfolded proteins was calculated by fitting the data to the Boltzmann sigmoidal equation in GraphPad Prism (version 8.0, GraphPad Software Inc., La Jolla, CA, USA).

### Cysteine residue reactivity assay

1Dx5-NTD^CAA^, 1Dx5-NTD^ACA^ and 1Dx5-NTD^AAC^ proteins were concentrated to above 3 mg/mL and reduced with 10 mM DTT for 30 min. DTT was removed by passing through the protein to a PD-10 column (GE Healthcare, Chicago, IL, U.S.) preequilibrated with 20 mM PBS pH 7.4. In a 25 μL protein solution, 2.5 μL of 10 mM potassium bromate was added to initiate the crosslinking. At time intervals of 1 min, 2 min, 4 min, 8 min, 30 min, 90 min, 180 min, 300 min, and 360 min, 0.5 μl of 100 mM iodoacetamide was added to terminate the reaction. Crosslinking products were analyzed with non-reducing SDS-PAGE described in the following, and the experiment was repeated in triplicates.

Non-reducing SDS-PAGE was performed where there was no DTT present in the loading buffer and molecular marker (Solarbio Life Sciences, Beijing, China) was loaded on the same gel as the reference. Gel containing a 4% stacking part and a 15% resolving part was electrophoresed at a constant voltage of 120 V with a Universal Electrophoresis Power Supply (WIX-EP600, WIX Technology Beijing Co., LTD., Beijing, China). When the bromophenol blue dye reached the bottom of the gel, the gel was stained with Coomassie Brilliant Blue to visualize protein bands. Gel images were captured with a SageCreation ChampGel^TM^ 5000 Automatic Gel Imaging and Analysis System (Beijing Sage Creation Science Co., LTD, Beijing, China) and analyzed with its related SageCapture^TM^ software with which the protein bands densities were calculated.

## Data Availability Statement

All data are available either in the article or in the Supplementary Information.

## Supporting Information

Supporting information containing Figure S1-8 and Table S1 are accompany this article.

## Acknowledgments

The authors are grateful to Prof. Chih-chen Wang, Prof. Lei Wang and Prof. Xi Wang (Institute of Biophysics, Chinese Academy of Sciences) for their support and encouragement, to Jifeng Wang (Laboratory of Proteomics, Institute of Biophysics, Chinese Academy of Sciences) for assistance in mass spectrometry, to Ms. Yun Feng (Center for Biological Imaging, Institute of Biophysics, Chinese Academy of Sciences) for help in microscopic imaging. DDY acknowledges support from the National Natural Science Foundation of China (32372372 and 31801482), and the National Laboratory of Biomacromolecules (2023kf02 and 2018kf09). DT acknowledges support from the National Science Foundation (CHE 2320256) and the Welch Foundation through the Collie-Welch Chair (F-0019).

## Author contributions

D.D.Y. and D.T. designed the research. B.X., J.F., J.G., Y.L. and Z.L. carried out the experiments. D.D.Y. and D.T. analyzed the data and wrote the manuscript.

## Notes

### Competing Interest Statement

The authors have declared no competing interest.

