## supplementary information for "Protein Folding accompanied by disulfide bond formation drives glutenin polymerization into multidimensional gluten networks"

This File Includes:

Figure S1-S7

Table S1

| | | | | | | |
|---|---|---|---|---|---|---|
| EGEASEQLQC | PGQ-WEE- | PTS-LQQ- | PGQ-GQQ-GQQ- | PGQ-WQQ- | PGQ-GQQ- | PTS-PQQ- |
| ERELQELQER | PEQ-GQQ-GYY- | SGQ-GQP-GYY- | VGQ-GQQ-AQQ- | PGQ-GQP-GYY- | PGQ-GQP-WYY- | PPQ-GQQ |
| ELKACQQVMD | PTS-PQQ- | PTS-LQQ- | PGQ-GQQ- | PTS-PLQ- | PTS-PQE- | LGQ-WLQ- |
| QQLRDISPEC | PGQ-LQQ- | LGQ-GQS-GYY- | PGQ-GQP-GYY- | PGQ-GQP-GYD- | SGQ-GQQ- | PGQ-GQQ-GYY- |
| HPVVVSPVAG | PAQ-GQQ | PTS-PQQ- | PTS-PQQ- | PTS-PQQ- | PGQ-WQQ- | PTS-LQQ- |
| QYEQQIVVPP | PGQ-GQQ-GQQ- | PGQ-GQQ- | SGQ-GQP-GYY- | PGQ-GQQ- | PGQ-GQP-GYY- | TGQ-GQQ- |
| KGGSFYPGET | PGQ-GQP-GYY- | PGQ-LQQ- | LTS-PQQ- | PGQ-LQQ- | LTF-SVA- | SGQ-GQQ-GYY- |
| TPPQQLQQRI | PTS-SQL- Q- | PAQ-GQQ- | SGQ-GQQ- | PAQ-GQQ-GQQ- | ART-GQQ-GYY- | SSYHVSVEHQ |
| FWGIPALLK | PGQ-LQQ- | PGQ-GQQ-GQQ- | PGQ-LQQ- | LAQ-GQQ-GQQ- | PTS-LQQ- | AASLKVAKAQ |
| -RYY- | PAQ-GQQ-GQQ- | PGQ-GQQ-GQQ- | SAQ-GQK-GQQ- | PAQ-VQQ-GQQ- | PGQ-GQQ- | QLAAQLPAMC |
| PSV-TC -PQQ- | PGQ-AQQ-GQQ- | PGQ-GQQ- | PGQ-GQQ- | PAQ-GQQ-GQQ- | PGQ-WQQ- | RLEGGDALSA |
| V -SYY- | PGQ-GQQ- | PGQ-GQP-GYY- | PGQ-GQQ-GQQ- | LGQ-GQQ-GQQ- | SGQ-GQH-WYY- | SQ |
| PGQ-AS -PQR- | PGQ-GQQ-GQQ- | PTS-PQQ- | PGQ-GQQ-GQQ- | PGQ-GQQ-GQQ- | PTS-PKL- | |
| PGQ-GQQ- | PGQ-GQQ- | SGQ-GQP-GYY- | PGQ-GQP-GYY- | PAQ-GQQ-GQQ- | SGQ-GQR- | |
| PGQ-GQQ-GYY- | PGQ-GQQ-GQQ- | PTS-SQQ- | PTS-PQQ- | PGQ-GQH-GQQ- | PGQ-WLQ- | |
| PTS-PQQ- | LGQ-GQQ-GYY- | PTQ-SQQ- | SGQ-GQQ- | PGQ-GQQ-GQQ- | PGQ-GQQ-GYY- | |

**Fig. S1** Amino acid sequence of glutenin 1Dx5. Blue represents the N-terminal domain and green corresponds to the C-terminal domain that are flanked the central repeat low complexity domain. Cystine residues C10, C25 and C40 are colored in orange. Hydrophobic amino acid residues Y52, F65, Y66 and W82 are colored in yellow.

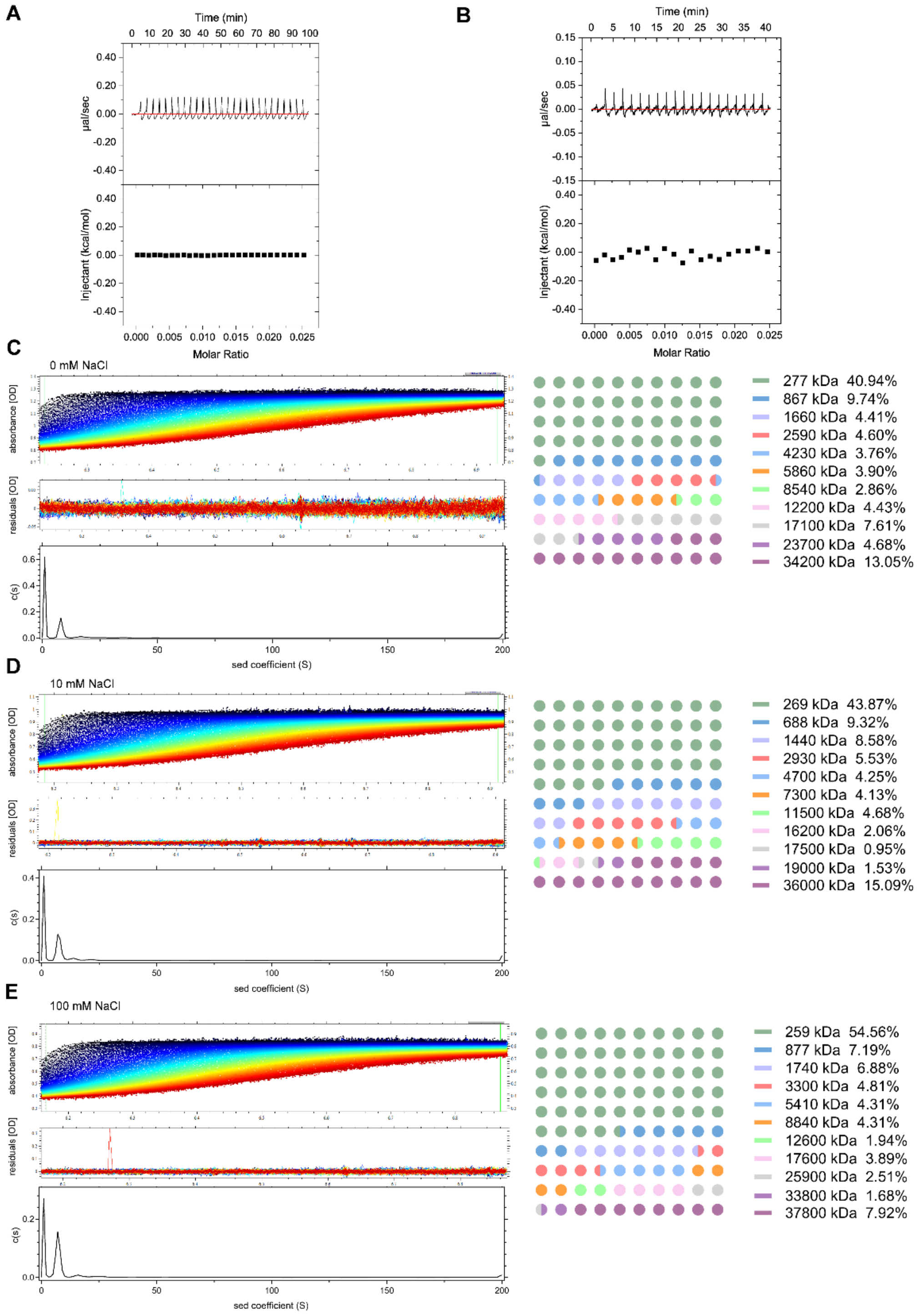


**Fig. S2** Additional characterization of 1Dx5-NTD association. **A-B**, Isothermal titration calorimetry thermographs of 200 μM cystine free 1Dx5-NTD$^{AAA}$ protein diluted into the PBS buffer which was used to dissolve the protein. Dilution was performed with a VP-iTC instrument (A) and repeated with an iTC$_{200}$ instrument with smaller volume (B). **C-E**, Sedimentation velocity profiles of cystine free 1Dx5-NTD$^{AAA}$ protein at different NaCl concentrations generated at 10k rpm with molecular weight distribution analyzed accordingly. c, 0mM; d, 10mM; and e 100mM NaCl.

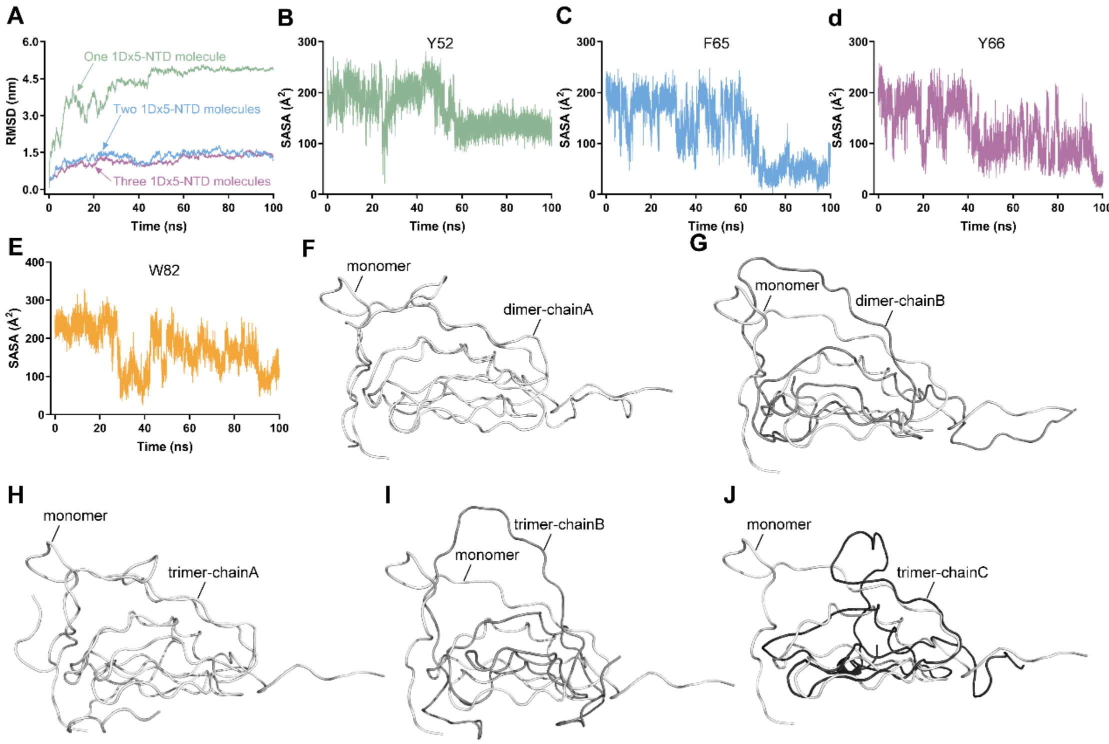


**Fig. S3** MD simulations of 1Dx5-NTD folding. **A**, Root-mean-square Deviation (RSMD) associated with folding of monomeric 1Dx5-NTD folding or interactions between two and three monomers other during the molecular dynamic simulations. **B**-**E**, Solvent accessible surface area (SASA) of the Y52 (b), F65 (c), Y66 (d), and W82 (e) side chains during 1Dx5-NTD monomeric folding. **F**-**G,** Superposition of 1Dx5-NTD onto itself and two interacting chains. **H-J,** Overlaid images of 1Dx5-NTD onto itself and three chains that interact with each other. Only two chains are displayed at a time in the trimer. There is structural heterogeneity in both the dimer and trimer.

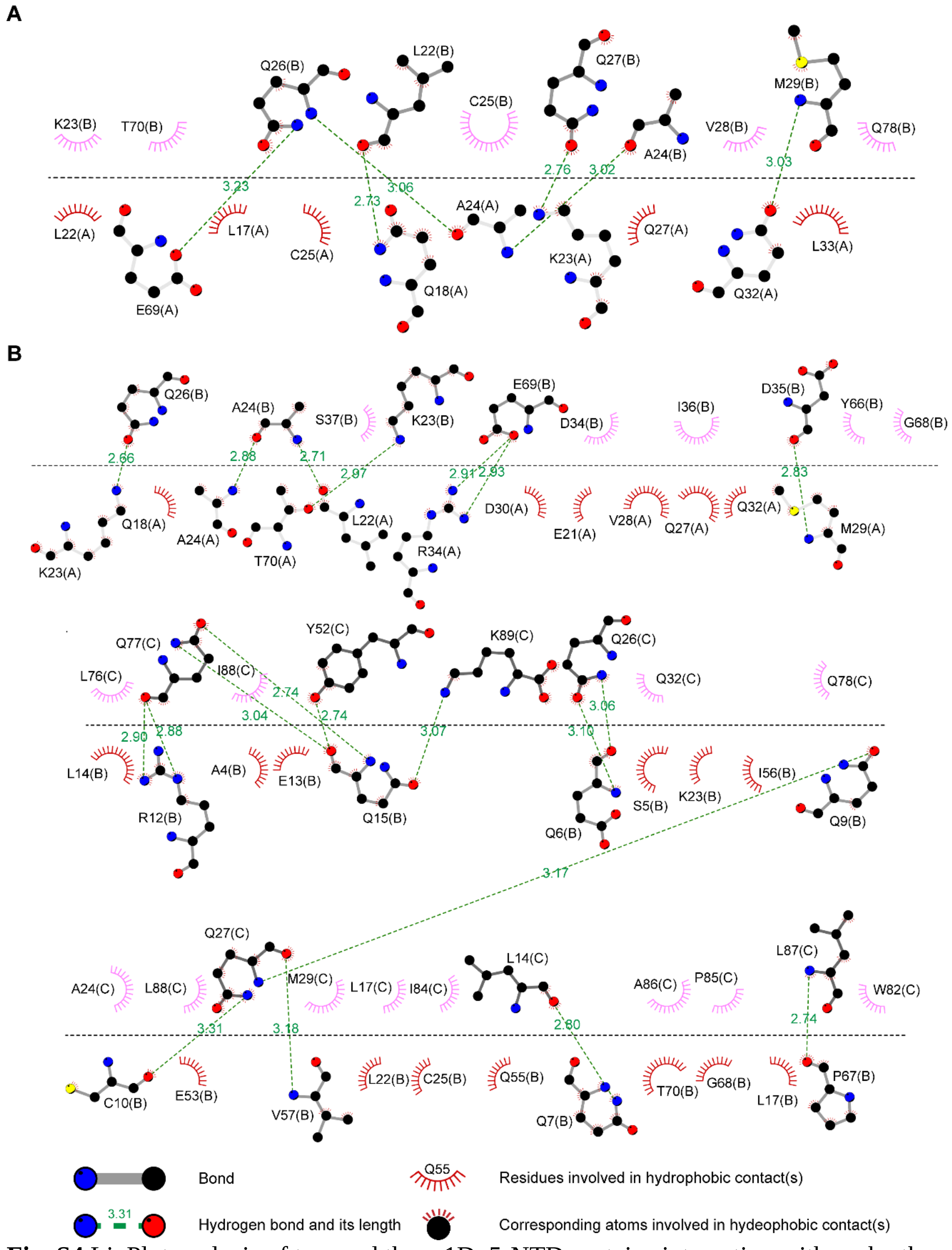


**Fig. S4** LigPlot analysis of two and three 1Dx5-NTD proteins interacting with each other. **A,** Details of two 1Dx5-NTD molecules interacting with each other. **B,** Details of three 1Dx5-NTD molecules interacting with each other. The horizontal dashed lines represent interface between two 1Dx5-NTD molecules.

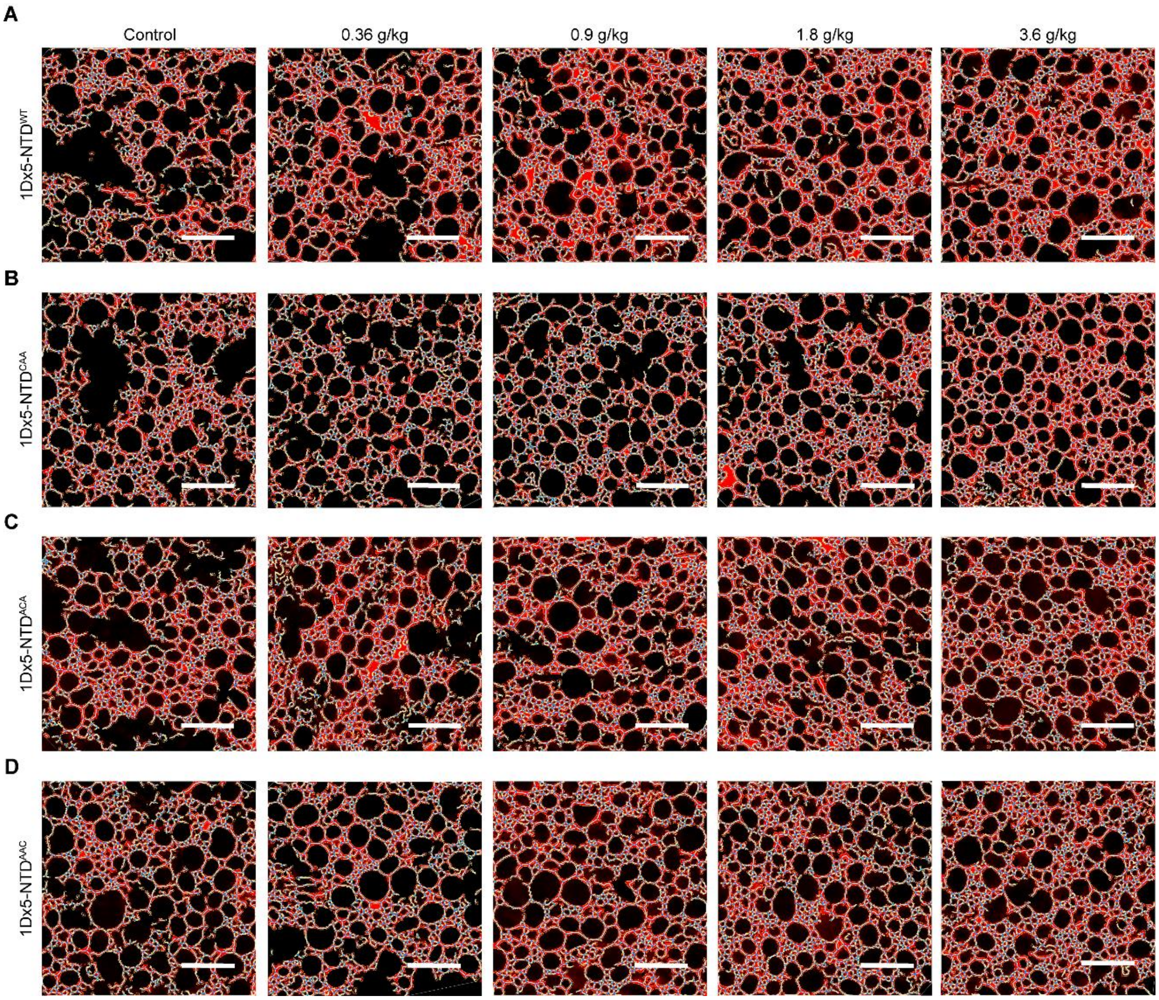


**Fig. S5** Confocal microscopic images of gluten network upon the addition of 1Dx5-NTD protein variants. **A**, Confocal microscopic images of gluten network upon the addition of wild type 1Dx5-NTD corresponding roughly to 10%, 25%, 50%, 100% of the total N-terminal domain of high molecular weight glutenin subunits. **B**-**D,** Images of gluten network upon addition of 1Dx5-NTD cystine mutants corresponding roughly to 10%, 25%, 50%, 100% of the total N-terminal domain of high molecular weight glutenin subunits. 1Dx5-NTD$^{CAA}$ denotes C25A/C40A mutation, 1Dx5-NTD$^{ACA}$ indicates C10A/C40A mutation, and 1Dx5-NTD$^{AAC}$ corresponds to C10A/C25A mutation. Red is the Rhodamine B-stained gluten network and blue lines are their analytical traces generated by the AngioTool software. White scale bar represents 50 μm.

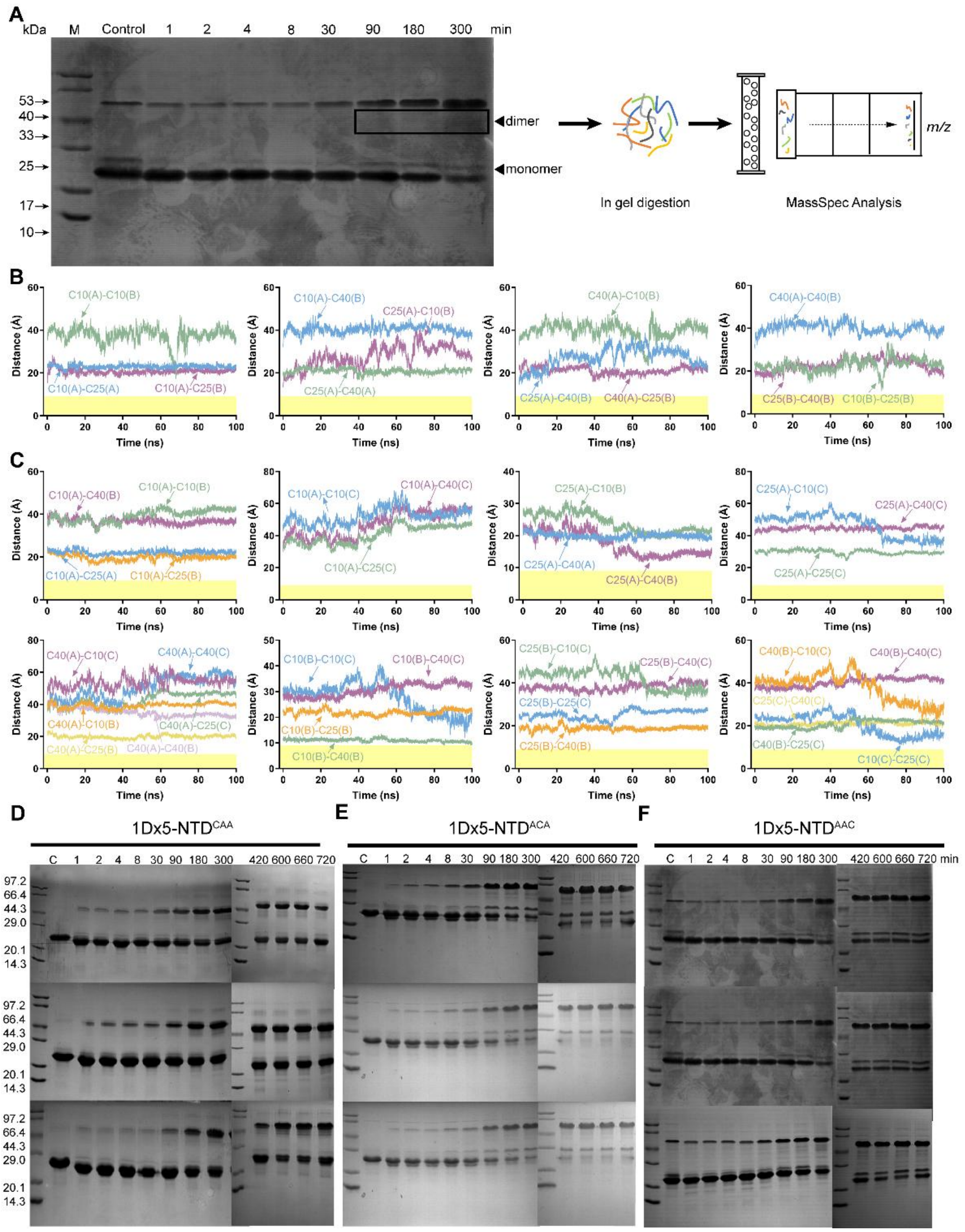


**Fig. S6** Crosslinking of 1Dx5-NTD. **A**, Non-reducing PAGE of 1Dx5-NTD crosslinking products at different time scale (1 min to 5 hours). The control was performed by oxidation under air. The band of two 1Dx5-NTD molecules crosslinked together were subjected to in gel protease digestion and MS/MS analysis. **B**, Distances between $C_{\alpha}$ of different cystines during two 1Dx5-NTD interaction. **C**, Distances between $C_{\alpha}$ of different cystines during three 1Dx5-NTD molecules interactions. **D-F**, Self-crosslinking of 1Dx5-NTD cystine mutant proteins. 1Dx5-NTD$^{CAA}$ corresponds to C25A/C45A mutation, 1Dx5-NTD$^{ACA}$ indicates C10A/C40A mutation, and 1Dx5-NTD$^{AAC}$ is the C10A/C25A mutation. The crosslinking products at 1, 2, 4, 8, 30, 90, 180, 300, 420, 600, 660, 720 min and the control group were analyzed on non-reducing SDS-PAGE. Numbers on the left are the molecular weights of the protein bands.

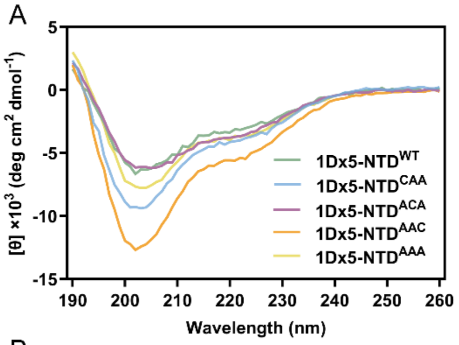


| Protein | α-helix (%) | β-sheet (%) | β-turn (%) | Random coil (%) |
|---|---|---|---|---|
| 1Dx5-NTD$^{WT}$ | 15.01 | 40.81 | 16.34 | 27.84 |
| 1Dx5-NTD$^{CAA}$ | 15.54 | 42.19 | 16.72 | 25.55 |
| 1Dx5-NTD$^{ACA}$ | 15.44 | 39.98 | 16.47 | 28.11 |
| 1Dx5-NTD$^{AAC}$ | 16.84 | 42.59 | 17.17 | 23.48 |
| 1Dx5-NTD$^{AAA}$ | 15.78 | 40.40 | 16.65 | 27.17 |

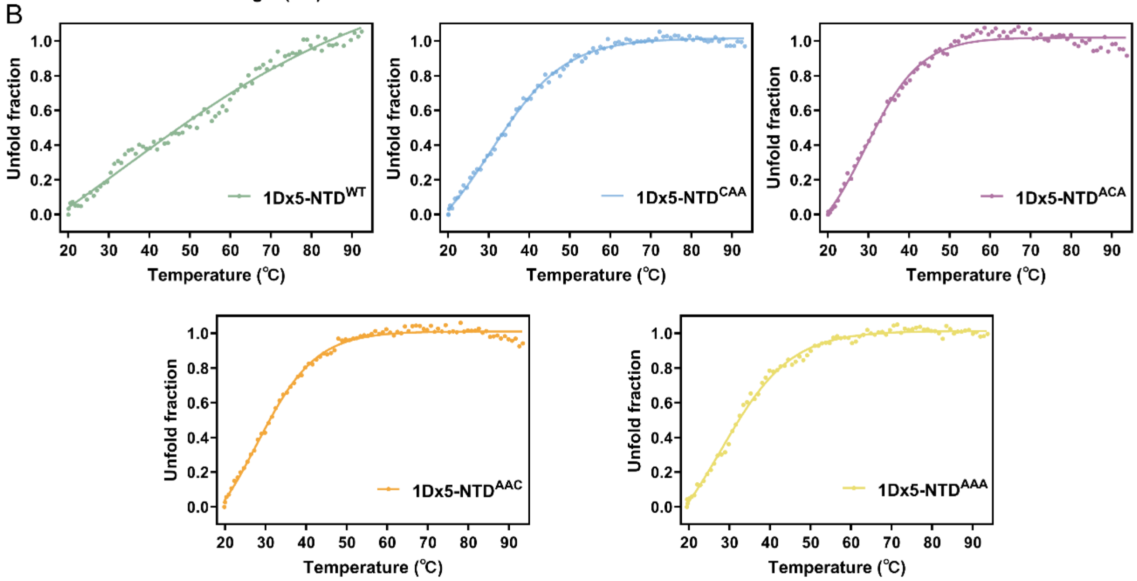


**Fig. S7** Impact of cystine residues on the secondary structure of 1Dx5-NTD. **A**, Circular dichroism spectrum of 1Dx5-NTD and cystine mutants. The secondary structure component of each protein variant was deducted from the CD spectrum and listed in the table. Mutations did not significantly alter their secondary structures. **B**, Melting curve of 1Dx5-NTD and cystine mutant. The CD signal of each protein at 222nm was recorded and the measurements were fit as solid curves. The melting temperature of each protein derived was not significantly different from each other.

**Supplementary Table 1 PCR Primers for Protein Variants Used in This Study**

| Protein Variant | Primer | Primer Sequences （5'-3'） | $^{*}T_m$ (℃) |
|---|---|---|---|
| 1Dx5-NTD$^{ACC}$ | C10A-Forward | TCTGAGCAACTACAGGCTGAGCGCGAG | 72.7 |
| | C10A-Reverse | GCCTGTAGTTGCTCAGAGGCCTCACCT | 72.7 |
| 1Dx5-NTD$^{CAC}$ | C25A-Forward | CGCGAGCTCAAGGCAGCCCAGCAGGTC | 77.8 |
| | C25A-Reverse | GCTGCCTTGAGCTCGCGCTCCTGGAGC | 77.2 |
| 1Dx5-NTD$^{CCA}$ | C40A-Forward | GACATTAGCCCCGAGGCCCACCCCGTC | 77.2 |
| | C40A-Reverse | GCCTCGGGGCTAATGTCTCGGAGCTGT | 74.7 |
| 1Dx5-NTD$^{Y52A}$ | Y52A-Forward | GGTCGCGGGACAAGCCGAGCAGCAAATC | 67.7 |
| | Y52A-Reverse | GATTTGCTGCTCGGCTTGTCCCGCGACC | 67.7 |
| 1Dx5-NTD$^{F65A}$ | F65A-Forward | GGCGGATCTGCCTACCCCGG | 59.9 |
| | F65A-Reverse | CCGGGGTAGGCAGATCCGCC | 59.9 |
| 1Dx5-NTD$^{Y66A}$ | Y66A-Forward | GCGGATCTTTCGCCCCCGGC | 59.9 |
| | Y66A-Reverse | GCCGGGGGCGAAAGATCCGC | 59.9 |
| 1Dx5-NTD$^{F65A}$ | F65A/Y66A-Forward | CGGATCTGCCGCCCCCGG | 57.9 |
| | F65A/Y66A-Reverse | CCGGGGGCGGCAGATCCG | 57.9 |
| 1Dx5-NTD$^{F81A}$ | F81A-Forward | CAACAACGTATAGCTTGGGGAATA | 53.4 |
| | F81A-Reverse | TATTCCCCAAGCTATACGTTGTTG | 53.4 |
| 1Dx5-NTD$^{W82A}$ | W82A-Forward | CCAACAACGTATATTTGCGGGAAT | 53.4 |
| | W82A-Reverse | ATTCCCGCAAATATACGTTGTTGG | 53.4 |

*Site directed mutagenesis was performed under the following conditions: 5 min at 94 ℃ for initial denaturation; 24 cycles of 30 s at 94 ℃, 30 s at each corresponding melting temperature, 6 min at 72 ℃; and a final elongation at 5 min at 72 ℃.
